# Reproducibility of neuroimaging studies of brain disorders with hundreds -not thousands- of participants

**DOI:** 10.1101/2022.07.05.498443

**Authors:** Ilan Libedinsky, Koen Helwegen, Udo Dannlowski, Alex Fornito, Jonathan Repple, Andrew Zalesky, Alzheimer’s Disease Neuroimaging Initiative, Alzheimer’s Disease Repository Without Borders Investigators, Michael Breakspear, Martijn P. van den Heuvel

## Abstract

An important current question in neuroimaging concerns the sample sizes required for producing reliable and reproducible results. Recent findings suggest that brain-wide association studies (BWAS) linking neuroimaging features with behavioural phenotypes in the general population are characterised by (very) weak effects and consequently need large samples sizes of 3000+ to lead to reproducible findings. A second, important goal in neuroimaging is to study brain structure and function under disease conditions, where effects are likely much larger. This difference in effect size is important. We show by means of power calculations and empirical analysis that neuroimaging studies in clinical populations need hundreds -and not necessarily thousands-of participants to lead to reproducible findings.

## Main Text

Finding reliable and reproducible effects is crucial for the progress of any scientific field. Key determinants in this pursuit are the quality of the methodology^1^, the number of participants in a study (i.e., sample size), and the magnitude of the studied effects (i.e., effect size)^2^. In a recent publication, Marek *et al*.^3^ provide important insights into the magnitude of effect sizes in neuroimaging MRI research and related sample sizes: brain-wide association studies (BWAS) linking neuroimaging features with behavioural phenotypes in the general population are characterised by (very) weak effects. Consequently, large sample sizes of over 3,000 participants are required to lead to reproducible effects. The largest effect sizes reported by Marek *et al*. concern correlations |r| as small as 0.06 to 0.16 across multiple imaging modalities and behavioural measures tested^3^.

Besides examining brain correlates of normative behaviour, another central goal in neuroimaging is to study brain structure and function under disease conditions, with the aim to shed light on pathophysiological mechanisms and identify potential markers of clinically relevant features^4^. Brain patterns related to disease conditions are likely more pronounced than general brain-behaviour associations; other factors remaining equal, smaller samples should be sufficient in these situations to obtain reliable and reproducible findings. This is important because clinical neuroimaging studies must balance reproducibility with cost-effectiveness and efficiency^2,5^. Here, we show by means of power calculations and empirical analysis that neuroimaging studies in clinical populations need hundreds -and not necessarily thousands-of participants to lead to reproducible findings.

Meta-and mega-analyses are a great tool to estimate effect sizes in disease conditions^5^. Meta-analysis of schizophrenia examining over 8,000^6^ participants report mean Cohen’s *d* effect sizes of -0.36 for case-control differences in grey matter volume (with effects ranging from -0.21 to -0.58 across brain regions)^6^. This is consistent with findings from the extensive studies by the ENIGMA consortium^7^, reporting group differences of *d*=-0.32 between schizophrenia patients and controls (*d*=-0.12 to -0.54 across 68 cortical regions, P<0.05 Bonferroni; 4,000+ patients and 5,000+ controls)^7^. While caution is needed when directly comparing effect sizes across study designs and methods^2^, these effect sizes are considerably larger than the effects found by Marek *et al*. for normative brain-behaviour associations^3^. In their study, the top 1% of all possible BWAS results reached |r|>0.06. In contrast, one of the smallest significant regional effects reported for schizophrenia^6^ is *d*=-0.21 which roughly converts^8^ to a |r|=0.11; the mean effect size of *d*=-0.32 as reported by ENIGMA^7^ matches^8^ a |r|=0.15.

This difference in effect size is important. Effect sizes are critical when determining the number of subjects required to obtain a desired level of statistical power (i.e. the probability of finding a true effect) in a study^2^. Power analysis for effects reported for schizophrenia (like a hypothesis driven region-of-interest group comparison of e.g. hippocampal volume, *d*=-0.43_6_) suggests samples of approximately 85 cases and 85 controls to be sufficient to reach the conventional standard of 80% power (two-sided t-test, *P*<0.05). In a BWAS, multiple brain regions are tested at the same time (for example, 333 regions are examined by Marek *et al*.) and correction for these multiple tests is needed, requiring larger sample sizes to maintain identical power^2^. Power simulations (Supplementary Methods; all code available) here indicate that 525 cases and 525 controls are needed to maintain 80% power for effect sizes as measured by ENIGMA^7^ when testing e.g. 333 regions (*P*<0.05 Bonferroni; Fig. 1a, solid lines; Supplementary Data 1). When alternatively testing less (100) or more (500) brain regions, samples of respectively 475 and 550 are needed per group (*P*<0.05 Bonferroni). These sample sizes can be smaller if a family-wise error correction is traded for a false discovery rate (FDR; Supplementary Data 1).

**Figure 1.**
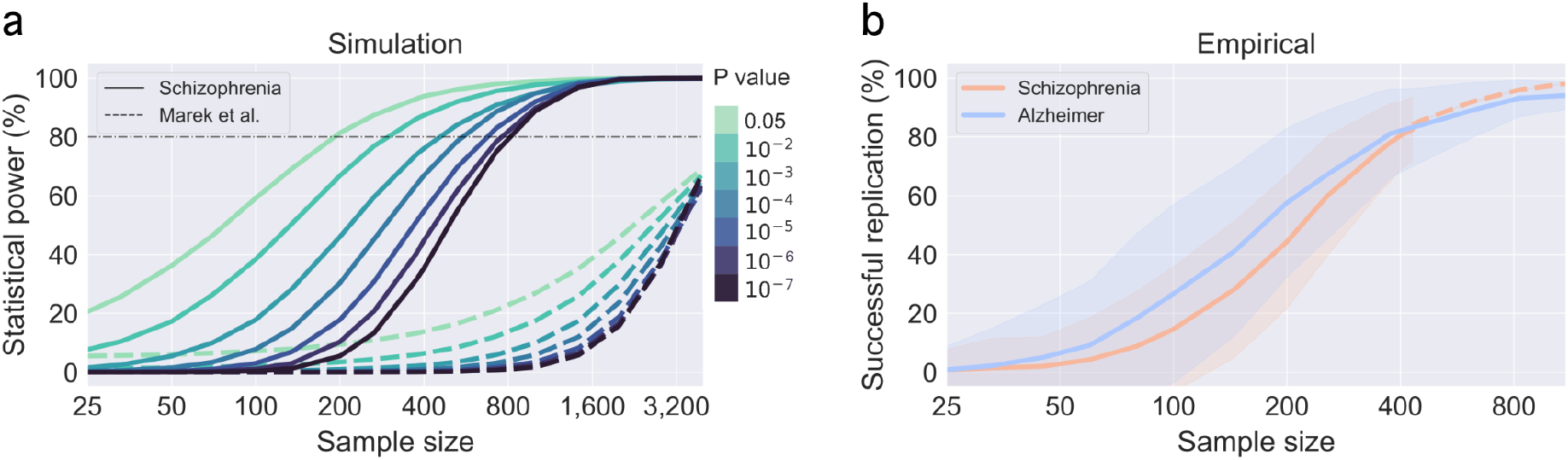
Power calculations and empirical analysis of replication rates. Panel **a** shows statistical power as a function of sample size and significance threshold (1,000 iterations). Solid lines depict power calculations based on ENIGMA^7^ effect sizes for schizophrenia (group differences in cortical thickness between patients and controls); for comparison, dashed lines show the power calculations reported by Marek *et al*.^3^ (Source Data Fig. 3 from the original article). Horizontal dash-dotted line corresponds to the field standard of 80% statistical power. Simulations indicate that sample sizes of 550 cases and 550 controls reach 80% statistical power for a BWAS with *P*<10^−4^; samples of 450 reach 80% power for a BWAS with *P*<10^−3^. Panel **b** shows replication rates for empirical data on schizophrenia (orange solid line) as a function of sample size (BWAS on 114 regions, tested against *P*<0.05 Bonferroni; 1,000 iterations; colored areas around the lines indicate one standard deviation above and below the mean across iterations). Orange dash line depicts extrapolated data (see Supplementary Methods). Empirical analysis shows that sample sizes of around 400 patients and 400 controls are needed to detect effects with a replication rate of 80%. Blue solid line shows replication rates for empirical data for Alzheimer’s disease (blue solid line). Sample size refers to the number of cases in each analysis with an equal amount of controls.

We further examined the replication of effects in real-world empirical data. We used anatomical T1-weighted data from five open neuroimaging cohorts to obtain cortical thickness data for 114 cortical areas in 866 schizophrenia patients and 1,897 healthy controls (all open data to foster replication; Supplementary Table 1 for demographics). We used the BWAS replication procedure as introduced by Marek *et al*. to assess the relationship between sample size and replication rate (Supplementary Methods). Two non-overlapping sets (“discovery” and “validation”) were randomly drawn and regional differences in cortical thickness were computed between patients and controls (two-sided t-test) across incrementally sample sizes. Iterating this process 1,000 times yielded the replication rate from the discovery to the validation set for each sample size. Using a strict *P* value threshold, a sample size of 400 patients and 400 controls is needed to achieve a replication rate of 80% (*P*<0.05 Bonferroni, 114 regions; Fig. 1b, orange line; Supplementary Data 2 shows BWAS results for alternative 68, 219 and 448 brain regions). Samples of approximately 200 are needed to reach a replication rate of 80% using FDR correction (*q*<0.05 FDR, 114 regions; Supplementary Data 2). The empirically derived effect sizes showed a mean Cohen’s *d* of -0.29, which closely matches the effect sizes reported by ENIGMA (*d*=-0.32)^7^ and used in the above power calculations.

We repeated the empirical analysis also for Alzheimer’s disease (2,297 patients and 2,267 controls; Supplementary Table 1). Samples of 400 patients and controls yield a replication rate of 80% (*P*<0.05 Bonferroni, 114 regions; Fig. 1b, blue line; Supplementary Data 2), and samples of 250 patients and controls reach replication rates of 80% using FDR (*q*<0.05; Supplementary Data 2).

The “reproducibility crisis”^9^ is a growing concern in many scientific fields, and neuroimaging is no exception^2,5,10^. Marek *et al*. importantly demonstrate that BWAS effect sizes of brain-behaviour relationships in the healthy population can be small, requiring large sample sizes to identify reproducible evidence. We agree with a general need for large sample sizes in our field, combined with more cross-validation, replication, and out-of-group analyses^11,12^. Care should however be taken when translating the Marek *et al*. results directly to other study designs (cross-sectional vs longitudinal designs) or populations (particularly patient samples; see several news^13^). Our simulation and empirical findings indicate that in clinical studies, when larger effect sizes are likely, sample sizes of hundreds -not thousands-are sufficient to detect reliable effects. We focused here on clinical cross-sectional studies, but effect sizes of similar magnitude are regularly reported for other phenotypes of major interest in our field, for example brain changes related to development or ageing (e.g. |r| from 0.1 to 0.5)^14^ and effects in clinical interventional studies (e.g. *d* from 0.1 to 0.6)^15^. While Marek *et al*. have not intended for their conclusions to be directly applied to other studies and populations, clarifying this important distinction is crucial to ensure that reproducibility, cost-effectiveness, and participant burden remain balanced across different contexts and to avoid erroneous extrapolations from normative to clinical research.

## Data availability

The datasets NUSDAST, MCIC, COBRE, BrainGluSchi, and NMorphCH are available from http://schizconnect.org. The CNP dataset is available from OpenNeuro (http://openneuro.org, accession number ds000030). The SCZIowa and B-SNIP datasets are obtained from the NIMH Data Archive (https://nda.nih.gov; NDAR ID: 2125 and 2274, respectively). SRPBS is available from https://bicr-resource.atr.jp/srpbs1600. The datasets ADNI1,2,3 are available from https://ida.loni.usc.edu. The datasets I-ADNI, EDSD, and ARWIBO are available from www.neugrid_2.eu. OASIS3 is available from www.oasis-brains.org. NACC is available from https://naccdata.org. Summary data of cortical thickness values from patients and controls for cortical regions are provided in Supplementary Table 2.

## Code availability

Code is available on https://github.com/dutchconnectomelab/brain-reproducibility

## Contributions

M.P.v.d.H. conceived the project. M.P.v.d.H, I.L. and K.H. analysed the data. M.P.v.d.H., I.L., and K.H. wrote the manuscript. A.Z, A.F., M.B., U.D., J.R. provided expertise and feedback on the text and analyses.

## Supplementary methods

### Dataset descriptions (summary)

#### Schizophrenia

High-resolution T1-weighted MRI data for schizophrenia was taken from the following public datasets (see also Supplementary Table 1 for an overview): *Northwestern University Schizophrenia Data and Software Tool* (NUSDAST, 136/182 patients and controls)^16^; *MIND Clinical Imaging Consortium* (MCIC, 29/25 patients and controls)^17^; *Centre for Biomedical Research Excellence* (COBRE, 70/81 patients and controls)^18^; *BrainGluSchi* (71/78 patients and controls)^19^; *Neuromorphometry by Computer Algorithm Chicago* (NMorphCH, 46/41 patients and controls)^20^; *Consortium for Neuropsychiatric Phenomics* (CNP, 50/125 patients and controls)^21^; *Phenomenology and Classification of Schizophrenia (Iowa Longitudinal Study)* referred as SCZIowa (84/121 patients and controls)^22^; *Bipolar & Schizophrenia Consortium for Parsing Intermediate Phenotypes* (B-SNIP, 239/401 patients and controls)^23^; and *Japanese Strategic Research Program for the Promotion of Brain Science* (SRPBS, 141/843 patients and controls)^24^. Subjects between 18 to 60 years old without any diagnosis or with a diagnosis of schizophrenia or schizoaffective-disorder were included in the analysis.

#### Alzheimer

High-resolution T1-weighted MRI data for patients with Alzheimer’s disease was taken from the following public datasets: *Alzheimer’s Disease Neuroimaging Initiative* (ADNI1, 520/205 patients and controls), ADNI2 (446/130 patients and controls), ADNI3 (149/92 patients and controls)^29^; *Italian Alzheimer’s Disease Neuroimaging Initiative* (I-ADNI, 222/0 patients and controls)^30^; *Open Access Series of Imaging Studies* (OASIS3, 174/713 patients and controls)^31^; *National Alzheimer’s Coordinating Center* (NACC, 116/322 patients and controls)^32^; *Alzheimer’s Disease Repository Without Borders* (ARWIBO, 403/632 patients and controls)^33^; *European Diffusion Tensor Imaging Study in Dementia* (EDSD, 269/173 patients and controls)^34^. Subjects between 50 to 95 years old without any diagnosis or with a diagnosis of probable Alzheimer’s disease or mild cognitive impairment were included in the analysis.

### Dataset descriptions (extended)

The *Northwestern University Schizophrenia Data and Software Tool* (NUSDAST) consist of neuroimaging data from patients with schizophrenia, healthy controls and their respective siblings^16^. Data was downloaded from XNAT central (http://central.xnat.org)^25^. The included data were high-resolution T1-weighted structural magnetization-prepared rapid gradient-echo (MPRAGE) scans collected on a 1.5T Siemens Vision scanner.

The *MIND Clinical Imaging Consortium* (MCIC) contains clinical and neuroimaging data from patients with schizophrenia and matched controls^17^. The MCIC project was supported by the Department of Energy under Award Number DE-FG02-08ER64581. MCIC is the result of efforts of co-investigators from University of Iowa, University of Minnesota, University of New Mexico, Massachusetts General Hospital. Data was downloaded from SchizConnect (http://schizconnect.org)^26^. T1-weighted high-resolution anatomical scans (MPRAGE) were acquired on a 1.5T Siemens Sonata and 3T Siemens Trio scanner.

The *Centre for Biomedical Research Excellence* (COBRE) includes multimodal neuroimaging data from patients with schizophrenia and matched controls^18^. Data was downloaded from SchizConnect (http://schizconnect.org)^26^. T1-weighted high-resolution anatomical scans (MPRAGE) were acquired on a 3T Siemens Triotim scanner.

The *BrainGluSchi* is a case-control study aiming to examine glutamate, glutamine and other neurometabolites in a sample of schizophrenia patients and healthy controls^19^. Data was downloaded from the SchizConnect (http://schizconnect.org)^26^. T1-weighted high-resolution anatomical scans (MPRAGE) were acquired on a 3T Siemens Triotim scanner.

The *Neuromorphometry by Computer Algorithm Chicago* (NMorphCH) is a longitudinal study collecting clinical, cognitive and neuroimaging data from schizophrenia and control subjects during two years^20^. Data was downloaded from SchizConnect (http://schizconnect.org)^26^. T1-weighted high-resolution anatomical scans (MPRAGE) were acquired on a 1.5T Siemens Vision scanner.

The *Consortium for Neuropsychiatric Phenomics* (CNP) examined brain function and anatomy for four psychiatric conditions, including comprehensive neuropsychological assessments^21^. Data was downloaded from OpenNeuro^27^ (http://openneuro.org, accession number ds000030). T1-weighted high-resolution anatomical scans (MPRAGE) were acquired on a 3T Siemens Trio scanner.

The *Phenomenology and Classification of Schizophrenia* (SCZIowa) consist of longitudinal data of symptoms, neuroimaging, and cognition from schizophrenia patients and controls^22^. Data was obtained from the NIMH Data Archive (NDAR ID: 2125). T1-weighted high-resolution anatomical scans (MPRAGE) were acquired on a 1.5T GE Sigma and 1.5T Siemens Avanto scanner.

The *Bipolar & Schizophrenia Consortium for Parsing Intermediate Phenotypes* (B-SNIP) aimed to examine a spectrum of clinical phenotypes within the schizophrenia-bipolar continuum, including 1st degree relatives and healthy controls^23^. Data was obtained from the NIMH Data Archive (NDAR ID: 2274). T1-weighted high-resolution anatomical scans (MPRAGE) were acquired on four 3T scanners (GE Signa, Philips Achieva, Siemens Allegra, and Siemens Trio).

The *Japanese Strategic Research Program for the Promotion of Brain Science* (SRPBS) includes multisite neuroimaging data of multiple psychiatric disorders (see https://bicr-resource.atr.jp)^24^. For the present study, the SRPBS Multi-disorder MRI Dataset (restricted) was included^28^. T1-weighted high-resolution anatomical scans (MPRAGE) were acquired on five 3T Siemens scanners (Verio, Spectra, VerioDot, TrimTrio and Trio) and two 3T GE scanners (Signa HDxt and MR750W).

The *Alzheimer’s Disease Neuroimaging Initiative* (ADNI1,2,3) database (adni.loni.usc.edu, for further information see www.adni-info.org^29^) was launched in 2003 as a public-private partnership, led by Principal Investigator Michael W. Weiner, MD. The ADNI study was approved by all Institutional Review Boards of all participating centres and all subjects provided written informed consent. The primary goal of ADNI has been to test whether serial MRI, positron emission tomography (PET), other biological markers, and clinical and neuropsychological assessment can be combined to measure the progression of mild cognitive impairment and early Alzheimer’s disease. Data from the first visit of each subject in the ADNI dataset were included. T1-weighted high-resolution anatomical scans (MPRAGE) were acquired on 3T Siemens Symphony, Siemens TrioTim and Siemens Prisma scanners.

The *Italian Alzheimer’s Disease Neuroimaging Initiative* (I-ADNI)^30^ is a cross-sectional study including patients with subjective memory impairment, mild cognitive impairment, Alzheimer’s dementia, and frontotemporal dementia. I-ADNI was provided by neuGRID (www.neugrid2.eu). T1-weighted high-resolution anatomical scans (MPRAGE) were acquired on 1.5T (GE Signa Excite, Siemens Avanto, Siemens Sonata, Philips Intera) and 3T (Philips Achieva, Siemens Allegra, GE Signa HDx) scanners.

The *Open Access Series of Imaging Studies* (OASIS) is a neuroimaging collection freely accessible to the scientific community^31^. OASIS3 is the latest release, consisting of a compilation of neuroimaging data of over a thousand participants (www.oasis-brains.org). T1-weighted high-resolution anatomical scans (MPRAGE) were acquired on 1.5T Siemens Vision and 3T Siemens TIM Trio scanners.

The *National Alzheimer’s Coordinating Center (*NACC)^32^ is prospective, standardized, and longitudinal clinical evaluation of the subjects in the National Institute on Aging’s ADRC Program. T1-weighted high-resolution anatomical scans were collected using 3T Siemens (Trio Tim, Verio, Allegra, Achieva, Sonata, Prisma, Skyra), Phillips (Gemini, Ingenuity) and GE (Genesis Signa, Signa HDx, Signa Excite, Signa, Discovery MR 750) scanners.

The *Alzheimer’s Disease Repository Without Borders* (ARWIBO) consist of comprehensive neuropsychological, EEG, neuroimaging, and biological data of patients with neurodegenerative diseases and healthy controls collected in over 10 years by a number of researchers of IRCCS Fatebenefratelli, Brescia, Italy^33,34^. The overall goal of ARWIBO is to contribute, thorough synergy with neuGRID (www.neugrid 2.eu), to global data sharing and analysis in order to develop effective therapies, prevention methods and a cure for Alzheimer and other neurodegenerative diseases. T1-weighted high-resolution anatomical scans (MPRAGE) were acquired on 1T Philips Gyroscan.

The *European Diffusion Tensor Imaging Study in Dementia* (EDSD) is a cross-sectional multisite dataset consisting of neuroimaging data from patients with Alzheimer’s Disease, mild cognitive impairment and healthy elderly controls^35^. EDSD was provided by neuGRID (www.neugrid2.eu). T1-weighted high-resolution anatomical scans (MPRAGE) were acquired on 1.5T (Siemens Sonata, Siemens Avanto) and 3T (GE Signa HDxt, Siemens Allegra, Siemens Trio, Siemens Achieva, Siemens TrioTim, Philips Intera) scanners.

### T1 data preprocessing

T1-weighted scans were processed using Freesurfer version v7.1.1^36^. Cortical thickness values were extracted for the Freesurfer cortical Desikan-Killiany brain atlas describing 68 cortical regions^37^ and two higher-resolution Cammoun^38^ sub-parcellations (describing 114, 219, and 448 regions). Data on the 114 brain atlas is reported in the main text; validation of results on the coarser 68 and more fine-detailed 219 and 448 atlases are described in Supplementary Data 2 and Supplementary Data 3. Data from subjects whose average cortical thickness across all regions exceeded three standard deviations above or below the study mean were excluded from further analysis. Effects of age, gender, and dataset were regressed out using linear regression analysis. Supplementary Table 1 provides a demographic overview of the included datasets in the analysis. Supplementary Table 2 provides summary statistics of cortical thickness values from patients and controls (within datasets) for each cortical brain region.

### Statistical power calculation

Effect sizes for schizophrenia were taken from the ENIGMA toolbox^39^ describing Cohen’s *d* effect sizes for 68 cortical regions as measured across 4,430 patients with schizophrenia and 5,057 controls (see ^40^ for details). Accompanied *P* value data available through the ENIGMA toolbox was used to identify at each *P* value threshold (*P*<0.05 to *P*<10^−7^) a disease map of brain regions showing a significant difference in cortical thickness between patient and controls. Power calculations were performed for a range of sample sizes (25 to 4,000, same as in Marek *et al*.^41^) and for a range of P values (from *P*<0.05 to *P*<10^−7^) as follows. First, simulated patient and control data was generated by drawing the given sample size from a standard normal distribution (mean=0, std=1). Next, disease effects were simulated in the patient group by adding region-wise disease effects to the patient population. Then, cortical thickness was compared between simulated patients and controls using a two-sided independent t-test. Across regions, power was calculated as the ratio between the real effects detected in the simulated sample and the total number of true positive effects present at the set *P* value threshold. The false negative rate was calculated as the ratio of real effects missed, and the false positive rate was computed by dividing the number of false positives by the number of true negative effects. Simulations were repeated 1,000 times at each *P* value threshold and for different sample sizes, with values averaged across iterations. Regional effect-sizes were resampled to respectively 114, 219, 448 values for power calculations for the other brain atlas resolutions.

### Empirical replication rates

Replication rates were computed following the procedure defined by Marek *et al*.^41^. Across 1,000 iterations, a “discovery” and a “validation” set of equal amounts of patients and controls were randomly drawn from the total pool of participants (non-overlapping samples between sets) for a range of sample sizes (15 logarithmic intervals, n= 25, 33, 45, 60, 80, 100, 145, 200, 256, 350, 400, 460, 615, 825, 1,100). Region-wise cortical thickness differences between patients and controls in the discovery set were tested using a two-sided independent t-test (results on 114 regions are reported in the main text, results for the alternative 68, 219, and 448 atlases are reported in Supplementary Data 2) for a range of *P* value thresholds (*P*<0.05 uncorrected, *q*<0.05 FDR, *P*<0.05/[number of regions] Bonferroni) across incrementally sample sizes. Next, for the validation set, brain features were similarly tested with a two-sided independent t-test between the drawn group of patients and controls. The replication rate was then quantified as the percentage of significant cortical regions in the discovery set that were similarly found significant in the validation set^41^. 1,000 iterations were performed for each sample size and statistical *P* value threshold, with percentages of replication rates averaged across iterations. Discovery and validation sets were fully independent in all iterations.

For the schizophrenia data, the largest subsample sizes that could be tested was 430 to achieve a split-half subdivision of the total set of 866 patients into two non-overlapping subsets. To provide an indication of statistical power for higher sample sizes (430 to 1,100), data was extrapolated by taking the mean and standard deviation of each dataset and region, sampling patient and control data points from the corresponding normal distribution. This procedure showed highly consistent predictions for power for the sample size range 25 to 430 (r=0.98, on average deviating 3.2 percentage points from the observed power).

#### Within-dataset subsampling

Replication rates were alternatively computed with subsampling by dataset. To this end, first, the discovery dataset and separate replication dataset were randomly selected from the total available cohorts, and subsequently the patient and control groups were sampled from within each of the cohorts. Consistent with the main analysis, sample size calculations for the schizophrenia effect sizes (114 brain regions) resulted in samples of 200 patients and 200 controls to reach 80% replication rate for FDR correction (*q*<0.05 FDR); a sample size of 230 (maximum sample size when pooling within-cohorts for schizophrenia) reached 54% replication rate when Bonferroni correction was applied (*P*<0.05 Bonferroni). Mean effect sizes were similar to the main analysis (mean Cohen’s *d*=-0.29). Replication rates were similar for the other brain parcellations (68, 219, and 448 brain regions). Sample size calculations for the Alzheimer data similarly showed samples of 256 subjects and 256 controls (114 brain regions) to reach a replication rate of 80% for FDR (*q*<0.05 FDR; Supplementary Data 3); and sample size of 400 patients and 400 controls to yield a replication rate of 80% for Bonferroni correction (*P*<0.05 Bonferroni, mean Cohen’s *d*=-0.25). Replication rates were consistent with other brain parcellations.

### Supplementary files

- Supplementary Table 1. Demographics of included participants in analyses (pdf)
- Supplementary Table 2. Cortical thickness values from patients and controls across regions for each dataset (xlsx)
- Supplementary Data 1. Summary data of power simulations across atlases (68, 114, 219, 448 regions) (xlsx)
- Supplementary Data 2. Summary data of empirical data across atlases (68, 114, 219, 448 regions) (xlsx)

## Acknowledgments

The research of M.P.v.d.H. was funded by a ERC Consolidator Grant (101001062) from the European Research Council (ERC).

Data used in preparation of this article were obtained from the NU Schizophrenia Data and Software Tool (NUSDAST) database (http://central.xnat.org/REST/projects/NUDataSharing) As such, the investigators within NUSDAST contributed to the design and implementation of NUSDAST and/or provided data but did not participate in analysis or writing of this report. Data collection and sharing for this project was funded by NIMH grant 1R01 MH084803.

MCIC data and demographic information was collected and shared by [University of Iowa, University of Minnesota, University of New Mexico, Massachusetts General Hospital] the Mind Research Network supported by the Department of Energy under Award Number DE-FG02-08ER64581.

COBRE data were downloaded from the Collaborative Informatics and Neuroimaging Suite Data Exchange tool (COINS; http://coins.mrn.org/dx), and data collection was performed at the Mind Research Network and funded by the Center of Biomedical Research Excellence (COBRE) grant 5P20RR021938/P20GM103472 from the NIH to V. Calhoun.

BrainGluSchi Data was downloaded from the COllaborative Informatics and Neuroimaging Suite Data Exchange tool (COINS; http://coins.mrn.org/dx) and data collection was funded by NIMH R01MH084898-01A1.

Data used in preparation of this article were obtained from the Neuromorphometry by Computer Algorithm Chicago (NMorphCH) dataset (http://nunda.northwestern.edu/nunda/data/projects/NMorphCH). As such, the investigators within NMorphCH contributed to the design and implementation of NMorphCH and/or provided data but did not participate in analysis or writing of this report. Data collection and sharing for this project was funded by NIMH grant R01MH056584.

Data and/or research tools used in the preparation of this manuscript were obtained from the National Institute of Mental Health (NIMH) Data Archive (NDA). NDA is a collaborative informatics system created by the National Institutes of Health to provide a national resource to support and accelerate research in mental health. Dataset identifier(s) are: 2125 and 2274. This manuscript reflects the views of the authors and may not reflect the opinions or views of the NIH or of the Submitters submitting original data to NDA.

Data collection and sharing for SRPBS was provided by the DecNef Department at the Advanced Telecommunication Research Institute International, Kyoto, Japan.

Data collection and sharing for this project was funded by the Alzheimer’s Disease Neuroimaging Initiative (ADNI) (National Institutes of Health Grant U01 AG024904) and DOD ADNI (Department of Defense award number W81XWH-12-2-0012). ADNI is funded by the National Institute on Aging, the National Institute of Biomedical Imaging and Bioengineering, and through generous contributions from the following: AbbVie, Alzheimer’s Association; Alzheimer’s Drug Discovery Foundation; Araclon Biotech; BioClinica, Inc.; Biogen; Bristol-Myers Squibb Company; CereSpir, Inc.; Cogstate; Eisai Inc.; Elan Pharmaceuticals, Inc.; Eli Lilly and Company; EuroImmun; F. Hoffmann-La Roche Ltd and its affiliated company Genentech, Inc.; Fujirebio; GE Healthcare; IXICO Ltd.;Janssen Alzheimer Immunotherapy Research & Development, LLC.; Johnson & Johnson Pharmaceutical Research & Development LLC.; Lumosity; Lundbeck; Merck & Co., Inc.;Meso Scale Diagnostics, LLC.; NeuroRx Research; Neurotrack Technologies; Novartis Pharmaceuticals Corporation; Pfizer Inc.; Piramal Imaging; Servier; Takeda Pharmaceutical Company; and Transition Therapeutics. The Canadian Institutes of Health Research is providing funds to support ADNI clinical sites in Canada. Private sector contributions are facilitated by the Foundation for the National Institutes of Health (www.fnih.org). The grantee organization is the Northern California Institute for Research and Education, and the study is coordinated by the Alzheimer’s Therapeutic Research Institute at the University of Southern California. ADNI data are disseminated by the Laboratory for NeuroImaging at the University of Southern California.

Data were provided [in part] by OASIS (T. Benzinger, D. Marcus, J. Morris; NIH P50 AG00561, P30 NS09857781, P01 AG026276, P01 AG003991, R01 AG043434, UL1 TR000448, R01 EB009352. AV-45 doses were provided by Avid Radiopharmaceuticals, a wholly owned subsidiary of Eli Lilly).

The NACC database is funded by NIA/NIH Grant U24 AG072122. NACC data are contributed by the NIA-funded ADCs: P30 AG019610 (PI Eric Reiman, MD), P30 AG013846 (PI Neil Kowall, MD), P50 AG008702 (PI Scott Small, MD), P50 AG025688 (PI Allan Levey, MD, PhD), P50 AG047266 (PI Todd Golde, MD, PhD), P30 AG010133 (PI Andrew Saykin, PsyD), P50 AG005146 (PI Marilyn Albert, PhD), P50 AG005134 (PI Bradley Hyman, MD, PhD), P50 AG016574 (PI Ronald Petersen, MD, PhD), P50 AG005138 (PI Mary Sano, PhD), P30 AG008051 (PI Thomas Wisniewski, MD), P30 AG013854 (PI Robert Vassar, PhD), P30 AG008017 (PI Jeffrey Kaye, MD), P30 AG010161 (PI David Bennett, MD), P50 AG047366 (PI Victor Henderson, MD, MS), P30 AG010129 (PI Charles DeCarli, MD), P50 AG016573 (PI Frank LaFerla, PhD), P50 AG005131 (PI James Brewer, MD, PhD), P50 AG023501 (PI Bruce Miller, MD), P30 AG035982 (PI Russell Swerdlow, MD), P30 AG028383 (PI Linda Van Eldik, PhD), P30 AG053760 (PI Henry Paulson, MD, PhD), P30 AG010124 (PI John Trojanowski, MD, PhD), P50 AG005133 (PI Oscar Lopez, MD), P50 AG005142 (PI Helena Chui, MD), P30 AG012300 (PI Roger Rosenberg, MD), P30 AG049638 (PI Suzanne Craft, PhD), P50 AG005136 (PI Thomas Grabowski, MD), P50 AG033514 (PI Sanjay Asthana, MD, FRCP), P50 AG005681 (PI John Morris, MD), P50 AG047270 (PI Stephen Strittmatter, MD, PhD).

Data collection and sharing of ARWIBO was supported by the Italian Ministry of Health, under the following grant agreements: Ricerca Corrente IRCCS Fatebenefratelli, Linea di Ricerca 2; Progetto Finalizzato Strategico 2000-2001 “Archivio normativo italiano di morfometria cerebrale con risonanza magnetica (età 40+)”; Progetto Finalizzato Strategico 2000-2001 “Decadimento cognitivo lieve non dementigeno: stadio preclinico di malattia di Alzheimer e demenza vascolare. Caratterizzazione clinica, strumentale, genetica e neurobiologica e sviluppo di criteri diagnostici utilizzabili nella realtà nazionale,”; Progetto Finalizzata 2002 “Sviluppo di indicatori di danno cerebrovascolare clinicamente significativo alla risonanza magnetica strutturale”; Progetto Fondazione CARIPLO 2005-2007 “Geni di suscettibilità per gli endofenotipi associati a malattie psichiatriche e dementigene”; “Fitness and Solidarietà”; and anonymous donors. For more information on the ARWIBO consortium can be found at www.arwibo.it.

We acknowledge the contribution of data made available through SchizConnect (http://schizconnect.org; funded by NIMH [grant 1U01 MH097435]), NIMH Data Archive (https://nda.nih.gov), OpenNeuro (http://openneuro.org), neuGRID (www.neugrid2.eu), and XNAT Central (http://central.xnat.org).

## References

1. Elliott, M. L. et al. What Is the Test-Retest Reliability of Common Task-Functional MRI Measures? New Empirical Evidence and a Meta-Analysis. Psychological Science 31, 792–806 (2020).

2. Zuo, XN., Xu, T. & Milham, M.P. Harnessing reliability for neuroscience research. Nat Hum Behav 3, 768–771 (2019).

3. Marek, S. et al. Reproducible brain-wide association studies require thousands of individuals. Nature 603, 654–660 (2022).

4. Woo, C.-W., Chang, L. J., Lindquist, M. A. & Wager, T. D. Building better biomarkers: brain models in translational neuroimaging. Nature Neuroscience 20, 365–377 (2017).

5. Button, K. S. et al. Power failure: why small sample size undermines the reliability of neuroscience. Nature Reviews Neuroscience 14, 365–376 (2013).

6. Haijma, S. V. et al. Brain Volumes in Schizophrenia: A Meta-Analysis in Over 18 000 Subjects. Schizophrenia Bulletin 39, 1129–1138 (2012).

7. van Erp, T. G. M. et al. Cortical Brain Abnormalities in 4474 Individuals With Schizophrenia and 5098 Control Subjects via the Enhancing Neuro Imaging Genetics Through Meta Analysis (ENIGMA) Consortium. Biological Psychiatry 84, 644–654 (2018).

8. Nakagawa, S. & Cuthill, I. C. Effect size, confidence interval and statistical significance: a practical guide for biologists. Biological Reviews 82, 591–605 (2007).

9. Baker, M. 1,500 scientists lift the lid on reproducibility. Nature 533, 452–454 (2016).

10. Ioannidis, J. P. A. Why Most Published Research Findings Are False. PLoS Medicine 2, e124 (2005).

11. Poldrack, R., Baker, C., Durnez, J. et al. Scanning the horizon: towards transparent and reproducible neuroimaging research. Nat Rev Neurosci 18, 115–126 (2017).

12. van den Heuvel, M.P., Sporns, O. A cross-disorder connectome landscape of brain dysconnectivity. Nat Rev Neurosci 20, 435–446 (2019).

13. https://www.nytimes.com/2022/04/19/magazine/mri-brain-activity-psychology.html; https://www.technologynetworks.com/neuroscience/news/stats-study-reveals-reason-for-replicability-crisis-in-neuroscience-359637;and https://www.nature.com/articles/d41586-022-00767-3

14. Lupton, M. K. et al. A prospective cohort study of prodromal Alzheimer’s disease: Prospective Imaging Study of Ageing: Genes, Brain and Behaviour (PISA). NeuroImage: Clinical 29, 102527 (2021).

15. de Vries, Y. A., Schoevers, R. A., Higgins, J. P. T., Munafò, M. R. & Bastiaansen, J. A. Statistical power in clinical trials of interventions for mood, anxiety, and psychotic disorders. Psychological Medicine, 1–8 (2022).

## Supplementary references

16. Wang, L. et al. Northwestern University Schizophrenia Data and Software Tool (NUSDAST). Frontiers in Neuroinformatics 7 (2013).

17. Gollub, R. L. et al. The MCIC Collection: A Shared Repository of Multi-Modal, Multi-Site Brain Image Data from a Clinical Investigation of Schizophrenia. Neuroinformatics 11, 367–388 (2013).

18. Aine, C. J. et al. Multimodal Neuroimaging in Schizophrenia: Description and Dissemination. Neuroinformatics 15, 343–364 (2017).

19. Bustillo, J. R. et al. Glutamatergic and Neuronal Dysfunction in Gray and White Matter: A Spectroscopic Imaging Study in a Large Schizophrenia Sample. Schizophrenia Bulletin 43, 611–619 (2017).

20. Alpert, K., Kogan, A., Parrish, T., Marcus, D. & Wang, L. The Northwestern University Neuroimaging Data Archive (NUNDA). NeuroImage 124, 1131–1136 (2016).

21. Poldrack, R., Congdon, E., Triplett, W. et al. A phenome-wide examination of neural and cognitive function. Sci Data 3, 160110 (2016).

22. Andreasen, N. C. et al. Progressive Brain Change in Schizophrenia: A Prospective Longitudinal Study of First-Episode Schizophrenia. Biological Psychiatry 70, 672–679 (2011).

23. Ivleva, E. I. et al. Gray Matter Volume as an Intermediate Phenotype for Psychosis: Bipolar-Schizophrenia Network on Intermediate Phenotypes (B-SNIP). American Journal of Psychiatry 170, 1285–1296 (2013).

24. Tanaka, S.C., Yamashita, A., Yahata, N. et al. A multi-site, multi-disorder resting-state magnetic resonance image database. Sci Data 8, 227 (2021).

25. Herrick, R. et al. XNAT Central: Open sourcing imaging research data. NeuroImage 124, 1093–1096 (2016).

26. Wang, L. et al. SchizConnect: Mediating neuroimaging databases on schizophrenia and related disorders for large-scale integration. NeuroImage 124, 1155–1167 (2016).

27. Markiewicz, C. J. et al. The OpenNeuro resource for sharing of neuroscience data. eLife 10 (2021).

28. Tanaka, S. C. et al. SRPBS Multi-disorder MRI Dataset (restricted). Synapse https://doi.org/10.7303/syn22317079 (2019).

29. Jack, C. R. et al. The Alzheimer’s disease neuroimaging initiative (ADNI): MRI methods. Journal of Magnetic Resonance Imaging 27, 685–691 (2008).

30. Cavedo, E. et al. The Italian Alzheimer’s Disease Neuroimaging Initiative (I-ADNI): Validation of Structural MR Imaging. Journal of Alzheimer’s Disease 40, 941–952 (2014).

31. LaMontagne, P. J. et al. OASIS-3: Longitudinal Neuroimaging, Clinical, and Cognitive Dataset for Normal Aging and Alzheimer Disease. (2019) doi:10.1101/2019.12.13.19014902.

32. Beekly, D. L. et al. The National Alzheimer’s Coordinating Center (NACC) Database: The Uniform Data Set. Alzheimer Disease & Associated Disorders 21, 249–258 (2007).

33. Frisoni, G. B. et al. Markers of Alzheimer’s disease in a population attending a memory clinic. Alzheimer’s & Dementia 5, 307–317 (2009).

34. Riello, R., Geroldi, C., Zanetti, O., Vergani, C. & Frisoni, G. B. Differential associations of Head and Body Symptoms with depression and physical comorbidity in patients with cognitive impairment. Int. J. Geriatr. Psychiatry 19, 209–215 (2004).

35. Brueggen, K. et al. The European DTI Study on Dementia — A multicenter DTI and MRI study on Alzheimer’s disease and Mild Cognitive Impairment. NeuroImage 144, 305–308 (2017).

36. Fischl, B. Automatically Parcellating the Human Cerebral Cortex. Cerebral Cortex 14, 11–22 (2004).

37. Desikan, R. S. et al. An automated labeling system for subdividing the human cerebral cortex on MRI scans into gyral based regions of interest. NeuroImage 31, 968–980 (2006).

38. Cammoun, L. et al. Mapping the human connectome at multiple scales with diffusion spectrum MRI. Journal of Neuroscience Methods 203, 386–397 (2012).

39. Larivière, S., Paquola, C., Park, By. et al. The ENIGMA Toolbox: multiscale neural contextualization of multisite neuroimaging datasets. Nat Methods 18, 698–700 (2021).

40. van Erp, T. G. M. et al. Cortical Brain Abnormalities in 4474 Individuals With Schizophrenia and 5098 Control Subjects via the Enhancing Neuro Imaging Genetics Through Meta Analysis (ENIGMA) Consortium. Biological Psychiatry 84, 644–654 (2018).

41. Marek, S. et al. Reproducible brain-wide association studies require thousands of individuals. Nature 603, 654–660 (2022).

42. Arslan, S. et al. Human brain mapping: A systematic comparison of parcellation methods for the human cerebral cortex. NeuroImage 170, 5–30 (2018).

